# Cell exit during EMT is mechanically triggered independently of E-Cadherin loss

**DOI:** 10.1101/2025.08.25.672137

**Authors:** Meritxell Font Noguera, Léa Roquin, Cyril Andrieu, Corinne Benassayag, Bruno Monier, Magali Suzanne

## Abstract

Epithelial-to-mesenchymal transition (EMT) is a fundamental cell process with essential functions in tissue and organ formation during development but also metastasis during cancer progression. The shift between these two states is driven by evolutionary conserved transcription factors. Cell exit from the epithelium of origin is considered to be a mere consequence of progressive adhesion loss due to transcriptional repression of genes encoding molecules such as E-Cadherin or cell polarity regulators. Here we show in the *Drosophila* leg that, counterintuitively, adhesions (as well as cell polarity) are maintained during cell exit, while an apico-basal force is applied on E-Cadherin complexes *in vivo*. By combining a genetic screen, optogenetics and laser ablation experiments, we found that this pulling force originates from transient basal filopodia-like protrusions, formed as soon as cells start to exit. Finally, apical cell detachment requires α-Spectrin activity, suggesting that the lateral cortex is required to efficiently transmit basal forces up to the apical cell pole. Our results indicate that, in a mature tissue, cell exit during EMT is a mechanically controlled process. Such a mechanism driving EMT without of E-Cadherin loss could fuel metastatic dissemination of collectively migrating cancer cells.

## INTRODUCTION

Epithelial-to-Mesenchymal Transition (EMT) is a cellular process which, by the conversion of static, adhesive epithelial cells into highly motile mesenchymal ones, is fundamental not only for embryonic development but also for tumor evolution ^1–3^. Epithelia are highly polarized with a marked apico-basal axis, basal adhesions such as Integrin receptors making the link to the underlying extracellular matrix (ECM) while intercellular adhesions are located apically ^4^. In particular, intercellular adhesion relies upon adherens junctions, E-Cadherin-dependent adhesive structures that mechanically connect cells of the tissue and make the link with the cytoskeleton ^5^. By opposition, mesenchymal cells have lost their apico-basal polarity and downregulated their adhesions ^2,6^. Degrees in shifting from an epithelial to a mesenchymal state are observed, including in tumors, leading to hybrid (or intermediate) states in which cells express both epithelial and mesenchymal molecular markers to a certain extent ^7,8^. In cancer, this eventually leads to metastatic dissemination from the primary tumor following EMT of single or small groups of circulating tumor cells, which is associated with poor clinical outcomes.

Extensive work in cell culture and model organisms has provided a rich picture of an evolutionary conserved regulatory network involving signaling pathways (e.g. TGF-β) and transcription factors (EMT-TFs such as SNAI1 in mouse) that govern decision making (i.e. to engage into EMT or not) in both development and tumorigenesis ^2,9^. Far less clear is our understanding of the execution of EMT, meaning not only how cells do remodel their shape and polarity but also how they detach from and move out of their epithelium of origin in order to become free to migrate in the underlying compartment, hence becoming ready to disseminate ^10^.

Repression of key adhesion and polarity proteins is a long-sought mechanism proposed to promote EMT. Indeed, many polarity proteins such as Disc Large (DLG), Lethal Giant Larvae or Crumbs (CRB) are transcriptional targets of EMT-TFs, while E-Cadherin is repressed in both invertebrates and vertebrates by most key EMT-TFs ^11–13^. The discovery more than two decades ago that Snail represses E-Cadherin transcription ^14,15^ coupled with the observation that E-Cadherin is lost in multiple cancer types (e.g. diffuse gastric cancers or lobular breast cancers) ^16^ and that downregulation of E-Cadherin is sufficient to promote EMT in cell culture while it facilitates metastasis after implantation in mice ^17^ prompted the idea of a long-term progressive loss of adhesion allowing eventually the detachment of the cell undergoing EMT. Yet, a growing number of arguments challenge this model. For instance, in some contexts, E-Cadherin is not transcriptionally repressed, as exemplified by endoderm formation in *Drosophila* ^18^. Furthermore, mouse embryos mutant for *Snail1* fail to downregulate *E-Cadherin* expression during mesoderm formation, yet many cells ingress ^19^. Finally, multiple types of mammary tumors rely upon E-Cadherin activity for efficient metastatic dissemination ^20^, consistent with the need for E-Cadherin during the migration of *Drosophila* endoderm cells ^21^.

If these works reveal that E-cadherin can, at least in some circumstances, be maintained through the whole EMT process, the strategies set up for cell exit without losing E-Cadherin remain elusive. Analyses of live embryos or explants have suggested that cell exit might be a quick phenomenon ^22,23^ and that it would rely upon actin remodeling downstream of Rho signaling ^24^. While E-Cadherin dynamics remain unexplored in these cases, these works suggest the existence of an active step driving apical cell detachment when cells initiate their exit from the epithelium.

To decipher how EMT cells could manage cell exit, we turned to EMT analysis in the *Drosophila* leg disc, a recent alternative *in vivo* inducible model. This model has been fruitful to reveal the existence of a transient apico-basal myosin II-dependent force which is concomitant to apical constriction at early EMT stages and which influences the morphogenesis of surrounding tissues ^25^. Here, we focused on later stages, when EMT cells exit the epithelium. We provide evidence that cells engaged in the EMT process maintain their apical adhesion and polarity. Interestingly, they transiently produce filopodia-like structures at their basal pole when they initiate cell exit, just prior apical detachment. These structures are associated with forces that are transmitted across the whole cell height. Finally, we show that the lateral α-Spectrin cortex is necessary to eventually detach apical adhesive complexes, freeing cells from their epithelium of origin. These results show that, during the EMT process, epithelial cell detachment is mechanically controlled and independent of E-Cadherin loss.

## RESULTS

### Uncoupling of force-associated cell exit from E-Cadherin and polarity loss during Snail-induced EMT

The *Drosophila* imaginal leg is a mature monolayer epithelium sitting on a basal membrane and forming a tube around a central lumen filled with extracellular matrix (**Fig.S1A-A’’**). We randomly expressed Snail in small groups of cells (i.e. clones, ∼5-10 cells) labelled by cytoplasmic staining (e.g. by expression of β-Galactosidase, β-Gal) in the fly leg, which led to EMT and eventually to the generation of mesenchymal-like cells (**Fig.1A**) ^25^. Of note, groups of EMT cells are not synchronized, as exemplified by mesenchymal-like cells in the proximity of still epithelial-like cells in **Fig.1A**. Similar clone behaviors were also observed in the wing disc (data not shown). This offers the opportunity to observe the whole sequence of events leading to cell exit in mature epithelial tissues.

**Figure 1.**
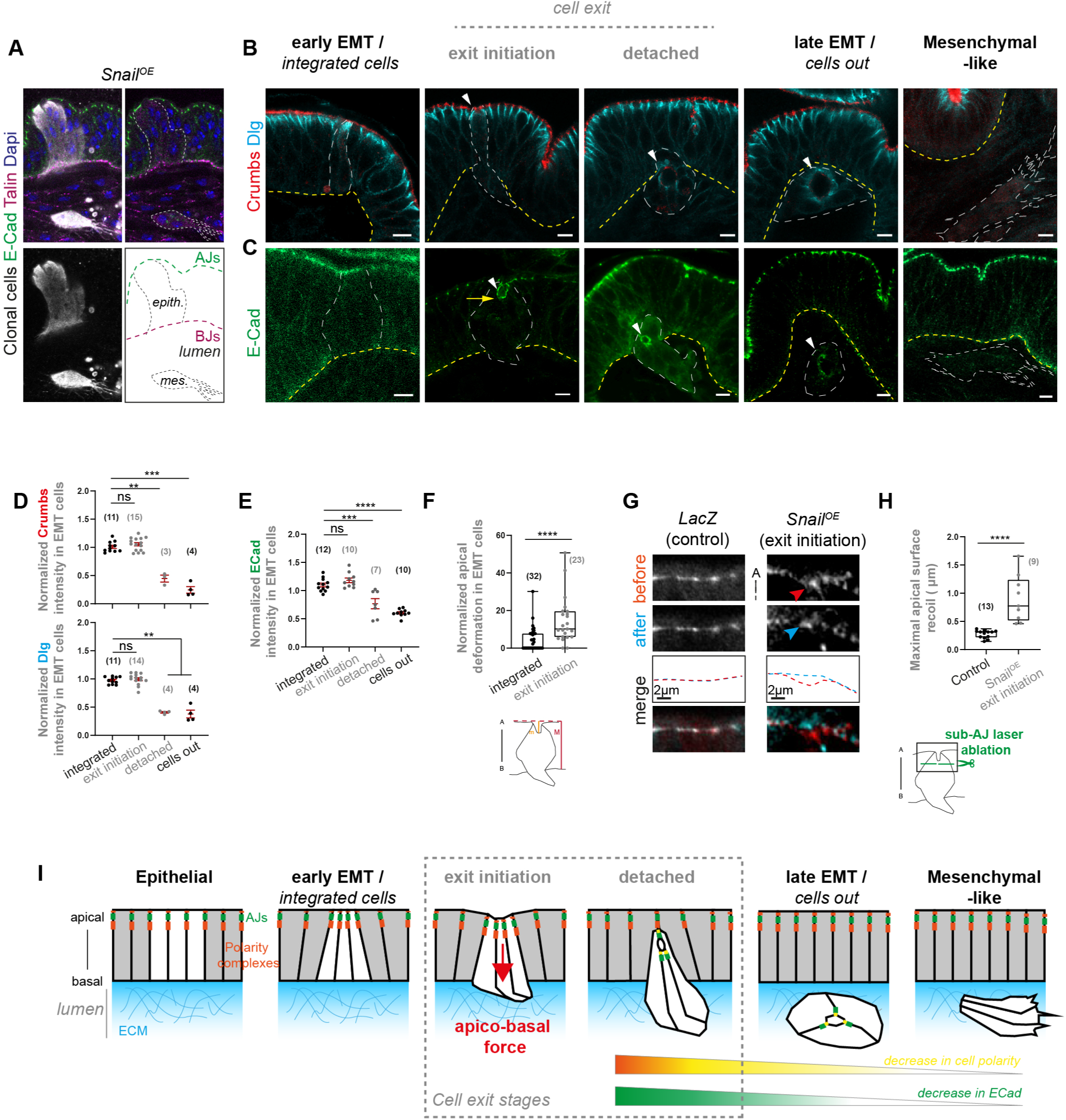
Uncoupling of force-associated cell exit from E-Cadherin and polarity loss during Snail-induced EMT. Related to Supplementary figure 1. (A) Snail-expressing clones (white) illustrating the two end-point of the EMT process in the *Drosophila* leg, with epithelial cells on top and mesenchymal-like in the luminal compartment (bottom). (B, C) Timeline of EMT stages ranging from fully integrated (early EMT stage, left) to cells completely out of the epithelium (late EMT stages), including cell exit (grey). Mesenchymal-like cells formed once EMT is completed are shown on the right. Outlines of Snail clones and basal of the epithelium are indicated by white and yellow dashed lines respectively, and clonal marker is shown in Supplementary Figure 1. Arrowheads indicate adhesive/polarity structures of Snail-cells observable until late stages, while the yellow arrow indicate apical surface deformation. (D, E) Quantifications of normalized intensity of polarity (D) and adherens junctions (E) markers along EMT progression based on (B, C). (F) Box plots illustrating apical deformation of the tissue ((m/M)*100)) at the level of EMT clones at early EMT (n=32) or exit initiation (n=23) stages. (G, H) Close up of transverse views (G) and associated box plots (H) of apical surface recoil after sub-apical photoablation in control (n=13) and EMT clones at cell exit (n=9). (I) Schematization of apico-basal force generation during cell exit preceding loss of cell polarity and of E-Cadherin. Scale bars: 5 µm unless indicated. In box plots, the black bar and whiskers indicate the median and the maximal range. In (D, E) the red bars indicate the mean and the standard error of the mean (SEM). n analyzed are indicated in brackets on the graphs. Statistical tests: Mann & Whitney in D, E, F and T-test in H. p-value: *, <0.05; **, < 0.005; ***, <0.001; ****, < 0.0001.

We first characterized the dynamics of apico-basal polarity and adherens junctions at the successive stages of conversion of epithelial cells into mesenchymal-like ones (**Fig.1B, C, Fig.S1B, C**). In this system, Snail EMT cells first undergo a series of mechanical steps at the onset of the EMT process, including an apical constriction (“early EMT” stage), consistent with apical constriction being one of the first event of the process *in vivo* ^10,26–28^. EMT cells then undergo cell exit, first by protruding their basal pole into the underlying luminal compartment (“exit initiation” stage), followed by apical detachment (“detached” stage). Once out in the lumen, EMT cells can transiently form cyst-like structures (“late EMT” stage) before acquiring migration features such as strong protrusions (“mesenchymal-like” stage). As expected, E-Cadherin, Crumbs and Dlg levels are nearly lost in mesenchymal-like cells (**Fig.1B, C**, right panels). Strikingly, polarity and E-Cadherin levels only start to drop once cells are apically detached and nearly out of the epithelium (**Fig.1D, E**). Cell exit is therefore initiated prior to any visible decrease in polarity or apical adhesion, supporting the fact that loss of adhesion or polarity are not the driver of cell detachment and that other mechanisms are at play in the *Drosophila* leg.

We wondered what could be the mechanism of apical detachment if apical intercellular junctions are maintained until cell exit. We observed that most EMT clones initiating cell exit (i.e. already invading the luminal compartment but still apically attached) present an indentation of the apical surface (**Fig.1C**, yellow arrow, quantification in **Fig.1F**). Apico-basal deformation of the apical surface have been previously associated with the existence of apico-basal forces ^25,29,30^. To directly test the presence of an apico-basal force when cell exit is initiated, we probed EMT cells by photoablation. Importantly, at the exit initiation stage, laser ablation of EMT cells just below the level of adherens junctions led to a basal-to-apical recoil of the apical surface, in contrary to similar ablations in control cells (**Fig.1G, H**). These results show that adherens junctions of EMT cells are subjected to apico-basal forces at the onset of cell exit.

Altogether, our results show that apical cell detachment, which is unlikely to be driven by loss of intercellular adhesion and polarity since it is initiated earlier, is associated with the existence of an apico-basal force which could drag EMT cells out of the tissue (**Fig.1I**).

### An in vivo screen identifies regulators necessary for efficient EMT

We next investigated which mechanism could be responsible of the generation of the apico-basal force associated with EMT cell detachment. Myosin II is the main molecular motor responsible of intra-cellular forces ^31^. Surprisingly, in most cases analyzed, we did not observe lateral apico-basal myosin II accumulations during cell exit (**Fig.S2A**), at a stage where apical accumulation is detected. This suggests that forces associated with EMT cell detachment are essentially myosin II-independent.

We speculated that the dynamics of cell shape changes during EMT and the generation of the apico-basal force at cell exit might rely upon actin cytoskeleton remodeling. One advantage of the leg model is that genes of interest can be manipulated specifically in EMT cells without targeting the whole epithelium. We therefore carried a genetic screen in which actin interactors or regulators were inactivated by RNA interference specifically in EMT cells, leaving unaffected epithelial cells of the environment. EMT cells, induced by expression of Snail, co-expressed a Histone-RFP transgene, in order to easily count them (**Fig.2A, Fig.S2B**). We screened about 200 RNAi lines and compared the number of Snail cells within the leg with the number of cells in which the neutral lacZ transgene was used to establish the baseline of EMT 48h after induction (**Fig.2B**). We expected that genes required for efficient EMT would lead to an accumulation of Snail cells that fail to exit the leg tissue following RNAi targeting. We identified 11 RNAi lines leading to a significant increase of EMT cells (2.5-fold, see **Fig.2B inset, Table S1**). Because an excess of cells after 48h could also be due to an increase in proliferation, we checked whether RNAi expression alone (i.e. in absence of Snail-induced EMT) could promote an increase in cell numbers. None of the RNAi lines tested in absence of Snail led to an increase in cell numbers (**Fig.S2C**), showing that their effect was specifically affecting EMT efficiency. Overall, we identified 11 genes potentially involved in Snail cells exit.

**Figure 2.**
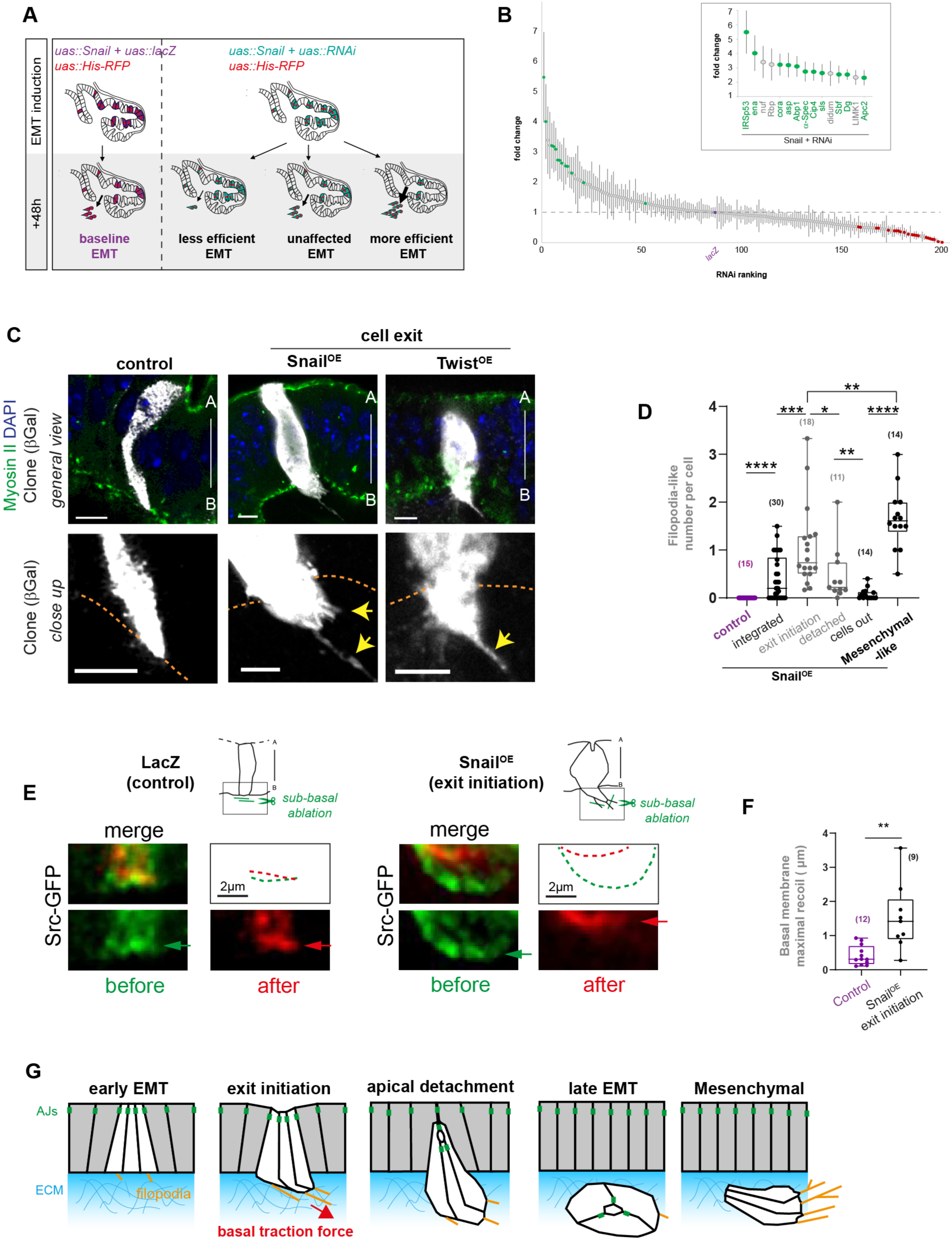
Identification of transient filopodia structures associated with basal traction forces during cell exit. Related to Supplementary Figure 2 and Supplementary Table 1. (A) Schematization of the genetic screen strategy. (B) Plot of the normalized number of Snail-cells (mean + SEM) after RNAi expression in the genetic screen. Green/red dots indicate significative RNAi compared to the neutral *lacZ* control (purple). Inset illustrates RNAi lines with maximal fold change. Related to Supplementary Table 1. (C) Images of cell exit induced by distinct EMT regulators, Snail and Twist. Clones are shown on the top and close up of the basal on the bottom. Arrows point at filopodia. The clone expressing Twist is at the initiation stage, the apical plane is shown in Fig.S2F. (D) Quantifications of basal filopodia numbers based on (C). (E, F) Close up transverse views (E) and associated box plots (F) of basal surface recoil after photoablation just outside the basal membrane in control (n=12) and EMT clones at cell exit (n=9). (G) Schematization of basal traction force and filopodia presence in EMT and mesenchymal-like cells. Scale bars: 5 µm unless indicated. n are indicated in brackets. In box plots, the black bar and whiskers indicate the median and the maximal range. Statistical tests: T-test in B; Mann & Whitney in D, F. p-values: *, <0.01; **, <0.001; ***, <0.0005; ****, <0.0001; ns, non significant.

### Transient filopodia-like protrusions associated with basal traction force prior apical detachment

In order to shed light into the dynamics of cells undergoing EMT, we used a Gene ontology (GO)-term analysis on the 11 candidate genes identified. While we retrieved coherent GO-terms, such as “cortical actin cytoskeleton”, we observed that the top 2 genes, *IRSp53* and *enabled (ena)* fell into an unexpected Biological function GO-term, “Filopodium assembly” (**Fig.S2D**). Although not associated with this GO-term, a third regulator of filopodia formation was retrieved: *Cip4*, the ortholog of TOCA-1 ^32^. In addition to lamellipodia, filopodia are associated with cell migration, including during cancer cell migration ^33,34^. Presence of filopodia was also reported in migrating cells once neural crest cells (NCC) have completed EMT ^10^. However, a role for filopodia in EMT cells before migration has not been documented as far as we know.

Filopodia are long, thin actin-rich membrane extensions, usually forming under the control of the RhoGTPase Cdc42 ^35,36^. Because they form transient, subtle structures, they can be easily overlooked. We therefore re-analysed the time-course of EMT events, staining again Snail EMT clones with cytoplasmic β-Galactosidase (β-Gal), but saturating the β-Gal channel in order to reveal potential membrane protrusions. Interestingly, we observed the presence of thin membrane protrusions in Snail cells from early stages of EMT (i.e. already in still fully integrated cells) onwards (**Fig.2C, D**). Filopodia-like protrusions generated by Snail cells are enriched in F-actin, as revealed by phalloidin staining, but devoid of myosin II, which can nevertheless accumulate at their base (**Fig.S2E**). Moreover, filopodia-like structures are also observed in EMT cells initiating cell exit (while still maintaining E-Cadherin) when an alternative EMT regulator, Twist, is induced in leg cells (**Fig.2C, Fig.S2F**). On the contrary to EMT cells, no membrane protrusions were observed in control clones (**Fig.2C, D**).

Importantly, both the number of filopodia *per* cell as well as the average filopodia length increase while Snail cells progress from early EMT to cell exit stages (**Fig.2D, Fig.S2G**). Filopodia numbers then drastically drop at late EMT stages when cell have managed to leave the epithelium, before strongly re-increasing as expected in mesenchymal-like, migrating clones (**Fig.2D**). Likewise, filopodia length tends to decrease between exit initiation and late EMT stages, before drastically re-increasing at the mesenchymal-like stage (**Fig.S2G**).

Of interest, filopodia have previously been associated with the existence of mechanical forces during tissue morphogenesis ^37,38^. Since we reported above the presence of forces at the exit initiation stage (**Fig.1G, H**), which is the timing of maximal filopodia presence in pre-migratory cells, we performed photoablation to test whether basal filopodia are engaged in force generation at that stage. Laser ablation just below the basal side of control epithelial cells does not produce marked recoil (**Fig.2E, F**), indicating that if cells are attached to their underlying extracellular matrix, they do not strongly pull on it. In sharp contrast, although in live tissues filopodia cannot be detected due to a lack in imaging resolution, laser ablation at the vicinity of the invading basal membrane of Snail cells at the exit initiation stage led to a stronger recoil of the basal membrane in most cases analyzed (**Fig.2E, F**). This strongly suggest that filopodia produced by Snail cells exert a pulling force on the ECM.

Altogether, our data indicate that the presence of filopodia-like structures during cell exit, a characteristic induced by two independent EMT regulators, is not a mere consequence of cell having started their migration, but rather a specific, transient feature of the cell exit program. It supports a model in which cells undergoing EMT while maintaining apical E-Cadherin are producing mechanical forces at their basal invading pole, likely *via* their transient protrusions adhering to the luminal ECM (**Fig.2G**).

### Cell exit is dependent of filopodia formation and ECM anchoring

We next sought to test for the contribution of the transient basal protrusion in favoring cell detachment and exit. Filopodia formation in Snail EMT cells is under the control of two molecules identified in the screen, IRSp53 and Ena, since their inactivation significantly reduces their number, albeit not completely (**Fig.3A, B**). Importantly, when these molecules are inactivated, EMT is slowed down (**Fig.3C**). Indeed, we quantified the repartition of clones integrated (*i.e.* early EMT) vs undergoing tissue exit at a given timepoint after induction of Snail expression in presence of normal (co-expressing *Luc^RNAi^*) or reduced filopodia numbers. We observed a shift towards early stages when filopodia are reduced due to *IRSp53* or *ena* RNAi expression (**Fig.3A, C**), indicating a slow-down or perturbation of the process when the formation of basal protrusions is impaired.

**Figure 3.**
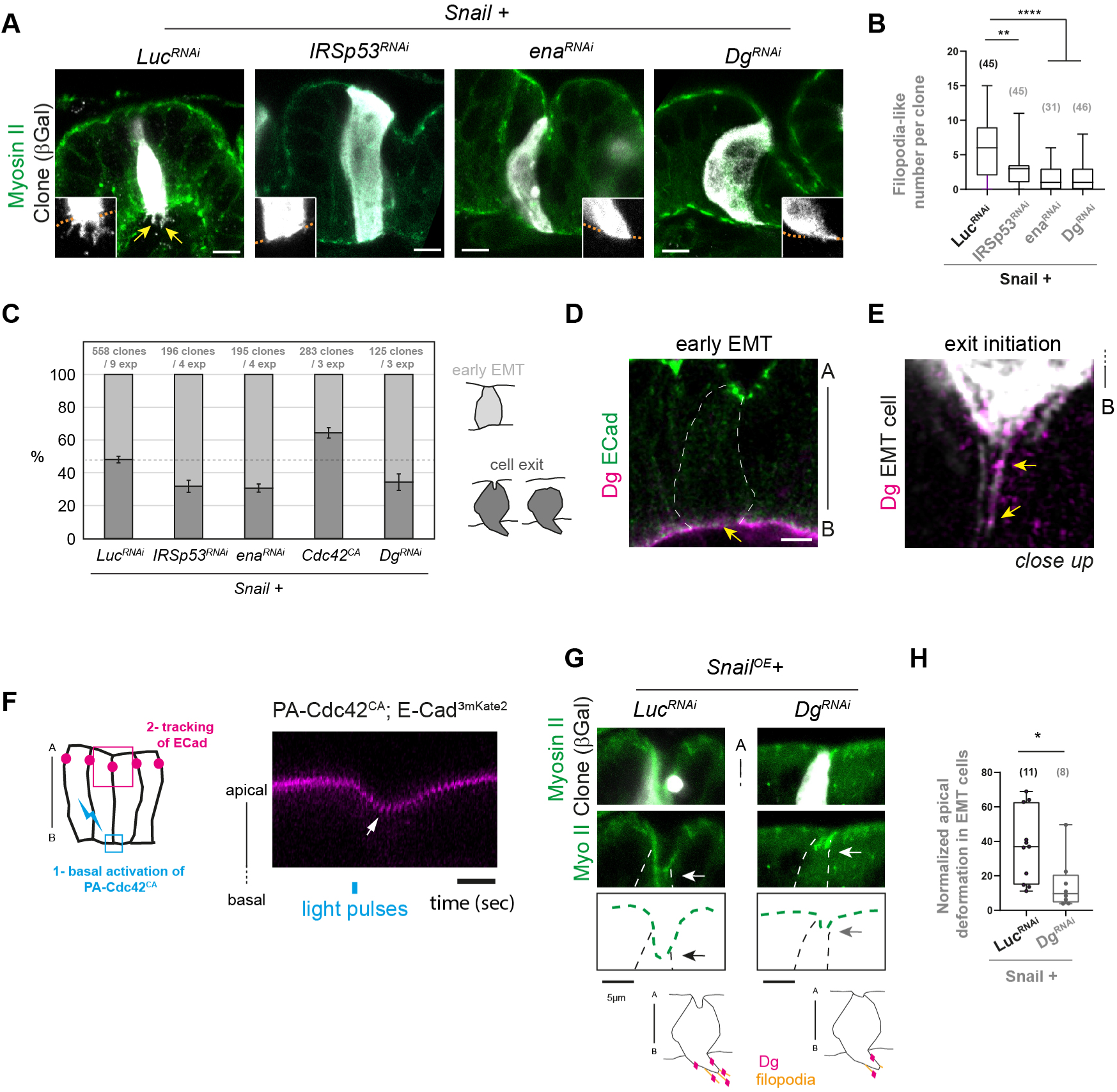
Filopodia-mediated EMT progression. Related to Supplementary Figure 3. (A, B) Illustrations of Snail-EMT clones and their close-up (A) and associated quantification of filopodia numbers (B) when co-expressing RNAi against *Luc* (control), *IRSp53*, *ena* or *Dg*. (C) Relative repartition of clones fully integrated (early EMT) or undergoing cell exit 42h after induction of Snail expression. Bars represent SEM. (D, E) Close up views of Dg in Snail cells at early EMT (D) and exit initiation (E) stages. Yellow arrows point at Dg staining respectively at the basal membrane (D) and filopodia (E). (F) Kymograph of the apical surface of non-EMT cells expressing the photoactivatable Cdc42CA construct. The blue line indicates the timing of photoactivation, the arrow points at the transient apicobasal displacement of E-Cadherin (n=7). (G, H) Illustrations of the apical surface (G) and associated quantification (H) of the apical deformation in Snail cells at the cell exit stage either co-expressing *Luc^RNAi^* (control, n=11) or *Dg^RNAi^* (n=8). Arrows indicate maximal apical surface deformation. Scale bars: 5 µm. In box plots, the black bar and whiskers indicate the median and the maximal range. n are indicated in brackets. Statistical tests: T-test in H, Mann & Whitney in B. p-values: *, <0.01; **, <0.001; ****, <0.0001.

We next wondered whether EMT could reciprocally be accelerated by increasing filopodia formation. As mentioned above, Cdc42 is a key regulator of filopodia formation ^35,36^, which is supported by the ability of its constitutive active form, Cdc42^CA^, to induce basal protrusion formation in the leg disc (**Fig.S3A**, see **Fig.2C** as a control). Interestingly, co-expression of Cdc42^CA^ with Snail is associated with an acceleration of EMT cell exit since less clones are at the early EMT stage at a similar timepoint after Snail induction (**Fig.3C**). Therefore, key regulators of filopodia structures are necessary and sufficient for EMT progression in pre-migratory cells, supporting a role for filopodia-like structures at timepoints clearly preceding the well-acknowledged migration of mesenchymal cells.

We reasoned that to play a role in the exit of EMT cells, filopodia-like protrusions might likely interact with the extracellular matrix. Coherently, we observed protrusions of EMT cells undergoing cell exit in close vicinity to the luminal ECM (**Fig.S3B**). Integrin complexes are unlikely to promote adhesion of filopodia to the ECM since their inactivation in the screen did not alter EMT efficiency (**Table S1**). In addition, both the mechanosensitive integrin binding partner Talin and the associated basal myosin II accumulation start to drop in Snail cells at early EMT stages, with Talin becoming barely detectable (**Fig.S3C-F**). However, another type of ECM receptor, Dystroglycan (Dg), was identified in the genetic screen (**Fig.2B**). Dg levels drop in early EMT cells, although it remains detectable (**Fig.3D**, **Fig.S3G, H**), reminiscent of basal adhesion regulation during early chicken gastrulation ^39^. Moreover, we observed a Dg punctiform staining in the filopodia-like protrusions (**Fig.3E**, arrows).

We next investigated the importance of Dg function for the EMT process. First, its loss-of-function strongly decreases the number of filopodia formed during EMT (**Fig.3A, B**). Second, downregulating *Dg* function also slows down EMT progression, similarly to *IRSp53* or *ena* depletion (**Fig.3A, C**). Our data suggest that basal EMT cell protrusions adhere to the ECM in the luminal compartment *via*, at least, the Dg transmembrane receptor, and that both protrusion formation/stability and EMT progression rely upon Dg activity.

### Basal-to-apical force transmission prior apical detachment

Could forces originating from the basal cell domain where filopodia form and Dg is localized be transmitted to the apical pole? To test this, we induced forces specifically in the basal domain using optogenetics. We expressed in non-EMT cells the PA-Cdc42^CA^ construct ^40^, a photoactivatable version of the filopodia inducer Cdc42^CA^. We then followed the behavior of adherens junctions labelled with E-Cadherin-3mKate. Strikingly, in these optogenetics experiments, a pulse of activation of Cdc42 on the basal cell side induced a transient apical-to-basal movement of adherens junctions (**Fig.3F**). This experiment shows that forces generated specifically at the basal cell side are transmitted up to adherens junctions.

We therefore tested whether filopodia forces produced at the basal cell pole are transmitted up to adherens junctions in EMT cells. We decreased Dg levels in EMT cells, thereby reducing filopodia numbers, and analyzed the impact on the apical surface deformation which is associated with the apico-basal force at the exit initiation stage. Importantly, the deformation of the apical surface is less intense in absence of Dg function (**Fig.3G, H**). Since Dg is specifically basal in EMT cells (**Fig.3D**), this indicates that forces originating at the basal pole impact the opposite, apical pole.

Overall, our results support the idea (i) that filopodia are mechanically-active structures in cells undergoing cell exit, likely due to their adhesion to the ECM via Dg-mediated adhesion, and (ii) that such forces are eventually transferred up to the levels of adherens junctions.

### Efficient apical detachment of EMT cells initiating cell exit requires the lateral α-Spectrin cortex

To challenge the notion of transmission from basal to apical, we looked for candidates among genes identified in the screen that could connect apical and basal cell poles. α-Spectrin proved to be a good candidate. Indeed, the Spectrin cytoskeleton links the actin cytoskeleton to the plasma membrane, where it provides structural stability ^41^, as is exemplified in C. elegans where the Spectrin cytoskeleton protects neurons from tearing during body movement ^42^.

We first characterized α-Spectrin dynamics during the sequential stages of EMT. In wild-type leg cells, endogenously GFP-tagged α-Spectrin is found at the basolateral cortex, up to adherens junctions, with a stronger enrichment towards the basal pole (**Fig.S4A, B**). Strikingly, α-Spectrin-GFP becomes depleted from the basal of Snail-cells at early EMT stages (**Fig.4A, Fig.S4C**). However, importantly, α-Spectrin remains localized to the lateral membranes, up to adherens junctions at any stage analysed up to late EMT stages (**Fig.4A**, yellow arrowheads). Its localization during cell exit is therefore consistent with a role in transmitting forces from the basal pole up to adherens junctions.

**Figure 4.**
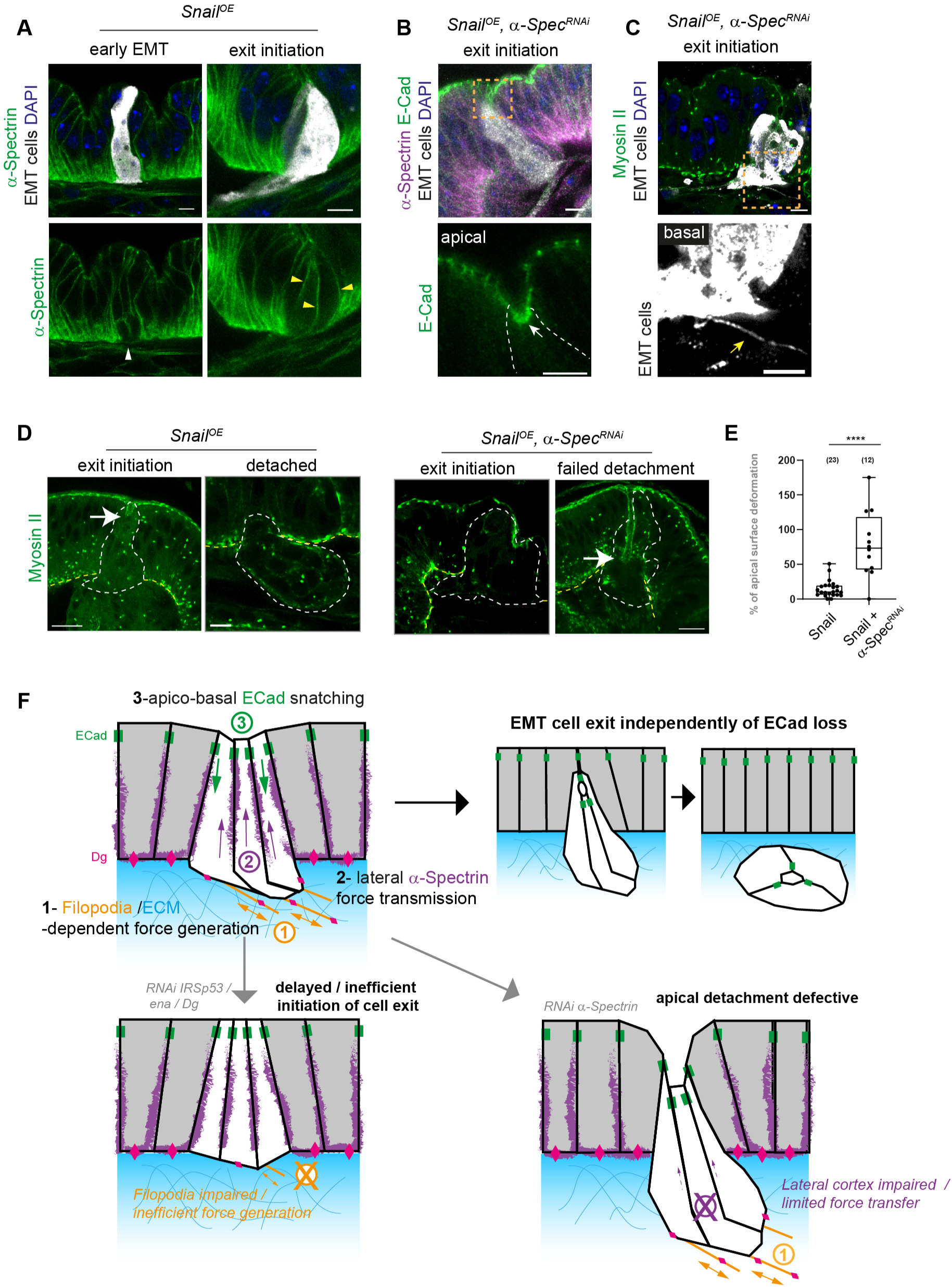
α-Spectrin is necessary for EMT-cells apical detachment. Related to Supplementary Figure 4. (A) Illustration of α-Spectrin distribution. White and yellow arrowheads point respectively at basal depletion and lateral maintenance. 9 clones analyzed in each context. (B, C) Snail cells undergoing cell exit despite α-Spectrin^RNAi^ depletion (top), with close up views (bottom) on apical (B) and basal (C) parts. White and yellow arrows point respectively at E-Cadherin maintenance and a filopodia. (D, E) Transverse views (D) and associated quantification (E) of EMT cells either control (left, n=23) or co-expressing α-*Spectrin^RNAi^*(right, n=12). Arrows point at maximal apical surface deformation. (F) A model of force generation and transmission driving cell exit in E-Cadherin-positive EMT cells. Global views of clones are provided in Supplementary Figure 4. Scale bars: 5 µm. In box plots, the black bar and whiskers indicate the median and the maximal range. Statistical tests: Mann & Whitney in E. p-values: ****, <0.0001.

We next asked whether depleting α-Spectrin from Snail-cells would impair cell exit. In sharp contrast with Snail cells depleted of *IRSp53*, *ena* or *Dg* which fail to efficiently invade the underlying luminal compartment, Snail cells depleted of *α-Spectrin* still do (**Fig.4B-D**), indicating that they are not stuck at early EMT stages. Consistently, we observed that α-Spectrin depleted Snail-cells maintain E-cadherin apically (**Fig.4B, Fig.S4D**), but still strongly reduce basal adhesion (using basal myosin II intensity as a proxy, **Fig.4D**), form filopodia (**Fig.4C, Fig.S4E**). Filopodia-dependent-basal forces could therefore still be generated when α-Spectrin is depleted. However, Snail cells depleted of α-Spectrin generated a stronger apical deformation than control Snail-cells (**Fig.4D**, arrows, quantification in **Fig4.E, Fig.S4F**), indicating that despite the basal force generated α-Spectrin depleted Snail-cells fail to efficiently detach from the apical of the epithelium. This failure in apical detachment could be a consequence of poor basal-to-apical force transmission in absence of the α-Spectrin cortex. These experiments highlight a critical role of the lateral cortex for completion of EMT cell exit in presence of adherens junctions.

## DISCUSSION

EMT is a key process, sustaining critical steps of development such as gastrulation as well as during pathological drift, such as fibrosis or cancer ^1,2,6^. In this latter pathology, EMT has been associated with acquisition of increasingly aggressive traits, including generation of cancer stem cells or increased resistance to drug treatments ^3,43^, and it is especially key in favoring cancer cell dissemination during metastasis. Providing a cellular basis and identifying molecular actors controlling this key step of cancer progression is therefore of prime importance. Based on the results obtained by combining high resolution imaging of small EMT clones, genetic screening and biophysical manipulation, we propose a sequential model of mechanical cell exit (**Fig.4F**): basal filopodia-like protrusions relying upon IRSp53, Ena and Dg activity would first generate traction forces at the basal cell pole. Basal forces would be transmitted laterally, at least in part via the α-Spectrin cortical skeleton, up to adherens junctions. Apico-basal forces, initially sufficient to displace more basally adherens junctions, would eventually reach a higher threshold at which E-Cadherin junctions would detach from wild-type neighbours, leaving EMT cells free to complete their exit from their epithelium of origin. Interestingly, forces applied upon adherens junctions could lead to opposite effects depending on their orientation: in *Drosophila*, forces applied parallel to adherens junctions (i.e. perpendicular to the cell membrane) stabilizes E-Cadherin complexes in mesodermal cells, while in ectodermal cells a shear stress applied perpendicular to E-Cadherin complexes (and parallel to the cell membrane) would destabilize them ^44,45^. Based on these results, we propose that apical constriction at early EMT stage could first stabilize adherens junctions between Snail cells, while subsequent filopodia-dependent apico-basal forces could create a destabilizing shear stress at the clone periphery.

Our results identify two types of filopodia-like structures: a first, transient one associated with the initiation of cell exit and preceding apical cell detachment, and a second, longer type associated with migration once cells have acquired a mesenchymal-like state. This indicates that filopodia associated with cell exit are not just a mere consequence of cells having started their migration but rather an intrinsic part of the program that drives cell exit, a program which precedes and can be uncoupled from the latter loss of apical E-Cadherin-dependent adhesion and of apico-basal polarity. Interestingly, the generation of filopodia associated with apical cell detachment is not a specific feature of Snail-induced EMT, since an independent EMT regulator, Twist, also triggers them. This suggests that basal filopodia (and their associated forces) could be a relatively general feature of EMT. Of interest, the presence of filopodia has previously been associated with cells that are converted from an epithelial (or endothelial) to a mesenchymal state in organisms as distinct as sea urchin, chicken or zebrafish ^46–48^. In sea urchin, these filopodia have even been associated with the generation of a force that impacts archenteron morphogenesis, while in *in sillico* modelling the presence of basal force-generating filopodia has been introduced to influence the directionality of cell exit ^47,49^. It would be of interest to re-examine the exact contribution of filopodia in these various models.

So far, a well-accepted sequence of EMT events suggests that polarity loss and transcriptional repression of apical adhesion components are early EMT events that would cause cell exit at later stages. Such a model is in accordance with the cases of E-Cadherin-negative cancers ^16^. Strikingly, dissecting the different steps of cell exit *in vivo* in *Drosophila*, we found that adhesion complexes are maintained during cell exit and only lost once cells are found in the underlying luminal compartment. This is of importance, as E-Cadherin is not systematically lost in cancers or during developmental EMT, providing cells with an unique opportunity to efficiently migrate collectively ^18,20,21^. It could also provide a mechanism explaining how circulating cancer cells can be generated at early tumor stages, for example in the pancreas prior to the appearance of “classical” signs of EMT (e.g. E-Cadherin loss, ZEB expression) ^50^. The precise characterization of the stages of exit of EMT cells in the *Drosophila* leg reported here will provide a framework for further investigations in mammalian systems and, hopefully at term help better control of metastasis.

## Acknowledgements

We thank Anne Pélissier-Monier for constructive comments on the manuscript. We thank Y. Bellaïche, K. Domsch, J. Kumar, M. Rolls, X. Wang and the stock centers BDSC and DGRC for sharing stocks and reagents. MS’s lab is supported by grants from the National Agency of Research (ANR, PRC AAPG2021, CellPhy), the Research Association against Cancer (ARC, Programme Labélisé AAP2020, ARCPGA12020010001154_1591) and the Medical Research Fundation (FRM, Equipes FRM, EQU202403018134).

## Author’s contributions

**Conceptualization**: B.M., C.B., M.S; **Methodology, Formal analysis and Investigation**: M.FN., L.R., C.A., B.M.; **Writing**: B.M., C.B., M.S; **Supervision**: B.M., C.B., M.S; **Resources**: B.M.; **Project administration**: M.S.; **Funding acquisition**: M.S.

## Materials and Methods

### Fly stocks & genetics

The individual identifier “BDSC_xxxx” is indicated for all stocks obtained from Bloomington Drosophila Stock Center (USA).

The sqh-eGFP^[^29^]^ knock-in line was used to reveal Myosin II distribution and was described previously in ^30^. hs::flp^[G5]^ is inserted in the attP3 site and was obtained from BDSC (BDSC_55817). The uas::Snail transgene was a gift from J. Kumar and previously used to induce EMT in the fly developing leg in ^25^. The act5C::FRT-CD2^stop^-FRT-Gal4, uas::Histone-RFP recombinant on the third chromosome has been isolated from BDSC stock (BDSC_51308).

For control experiments, a uas::lacZ transgene was used, either inserted on the third chromosome (BDSC_8530) or on the second chromosome uas::lacZ^[j]^. This latter stock was obtained by backcrossing the (BDSC_8529) stock in a *white* background. Alternatively, an uas::RNAi against Luciferase (BDSC_31603) was used as a control line.

uas::RNAi lines were obtained essentially from the Bloomington Drosophila Stock Center (BDSC), unless indicated. The list of the uas::RNAi lines used in the screen is indicated in Supplementary Table 1. Additional uas lines: uas::Cdc42[CA] (BDSC_4854) and uas::Twist-HA (gift from Katrin Domsch, Germany)

Additional fluorescent molecules tagged at their endogenous locus are E-Cadherin-GFP^[KI]^ (BDSC_60584), Talin-eGFP[KI] (this study, see below), Talin-mCherry^[MI00296]^/TM6B (BDSC_39648), Crumbs-GFP A[KI] (BDSC_99495), E-Cad-3mKate2[KI] ^51^, Dg-GFP[KI] ^52^, Vkg-TagRFPt[KI] ^53^, Vkg-GFP[KM502] (DGRC_109849) and α-Spectrin-GFP^[WeeP386]^. The α-Spectrin-GFP^[WeeP386]^ line is a gift from Melissa Rolls (The Pennsylvania State University, USA). It was described as a GFP insertion at the endogenous locus, between amino acids 1330 and 1331 (repeat 12) ^54^, which corresponds to an insertion after the first coding exon. The α-Spectrin-GFP line is homozygous viable. In the fly leg, it exhibits a pattern similar to the pattern of the protein detected with the 3A9 monoclonal antibody (from DSHB, USA). One difference however is that the GFP-trap protein is enriched at adherens junctions, a structure not labelled by the 3A9 antibody. Talin-eGFP[KI] is an homozygous viable knock-in line designed and generated by homologous recombination by InDroso functional genomics (Rennes, France). The resulting flies were validated by sequencing.

For laser ablation experiments, Snail-expressing cells were detected by co-expressing an uas::fluorescent transgene: uas::tdTomato (BDSC_92756), uas::Src^1-89^-GFP (BDSC_26157), uas::2xeGFP (BDSC_ 6874).

For optogenetics experiments, we used an uas::PA-Cdc42[CA] construct ^40^. Expression was achieved in the distal leg using the ap::Gal4^[md544]^ driver (BDSC_3041), combined with E-Cad-3mKate2^[KI]^.

### Random EMT induction in imaginal discs

For the characterisation of EMT dynamics and the function of the genes identified in the screen, we randomly induced small clones. The hs::flp^[G5]^ flippase was used either in presence of sqh-eGFP^[KI],^ as in the screen, or in absence of it to control the activity of the act5C::FRT-CD2^stop^-FRT-Gal4 transgene (BDSC_4780) and eventually drive expression of “uas” transgenes. The system was activated by a short heat shock pulse (10-12’) at 37°C. Crosses were then transferred back at 25°C for further development. Prepupae (white pupae (WP)/ WP+3 hours or eventually third instar larvae) were then dissected for fixation or live tissue analysis 42h after heat shock (unless an alternative timing is indicated).

Clones of interest were identified thanks to the expression of neutral markers (lacZ, tdTomato, cytoplasmic or membrane-bound GFP). EMT was driven by expressing *uas::Snail* or alternatively *uas::Twist* transgenes. A *uas::lacZ* transgene was used instead as a control. When uas::RNAi were co-expressed with uas::Snail, the neutral uas::RNAi Luc transgene was co-expressed instead to produce “control EMT”.

### EMT genetic screen methodology

For the genetic screen, a stock containing the following alleles and transgenes was crossed to individual uas::RNAi lines, or uas::lacZ as a control : w, sqh-eGFP^[KI]^, hs::flp^[G5]^; uas::Snail^FL^; act5C::FRT-CD2^stop^-FRT-Gal4, uas::Histone-RFP. X chromosome was homozygous, second and third chromosomes were maintained over a SM5-TM6B, Tb attached balancer. Crosses were done at 25°C.

Snail expressing cells systematically co-expressed an uas::Histone-RFP transgene which allowed their recognition in the leg. An RNAi targeting an actin related molecule was specifically expressed in Snail cells and then the efficiency of these cells to delaminate from the leg disc was assessed. Induction was done by heat shocking larvae for 12’ at 37°C. Larvae were transferred back at 25°C until dissection.

Around 10-15 leg disc per RNAi and EMT control condition were dissected after 48h of clones’ induction, fixed and imaged with confocal microscopy. Sqh-GFP and DAPI staining were used to counterstain the whole leg disc. To assess EMT efficiency, the number of Snail nuclei, indicator of Snail cell number in the leg, was counted in 3D legs by using the spot function of Imaris software. The mean number of Snail cells in the leg disc for every RNAi condition was normalized by their associated EMT control heat-shocked during the same cross series. The ranking of Snail cells number in the leg expressing each RNAi is indicated in Table 1. GO Enrichment Analysis was performed in the Gene Ontology website (https://geneontology.org/). For the secondary screen, we used the same strategy, except that the uas::Snail transgene was replaced by the uas::lacZ^[J]^ transgene in the testerline.

### Immunostainings

Leg discs were dissected at the early prepupal stage (white pupa-white pupa+3h), except for the screen in which both late 3^rd^ instar larvae and early prepupae (before peripodial membrane removal) were dissected. Dissection were carried out in PBS 1x. Tissue were fixed by paraformaldehyde 4% diluted in PBS 1x during 20 minutes. Then the samples were washed and saturated in PBS 1x, 0.3% triton x-100 and BSA 1% (BBT). Next, the samples were incubated overnight at 4°C with primary antibodies diluted in BBT. Samples were washed for 1h in BBT before a 2h incubation at room temperature with secondary antibodies diluted in BBT. Finally, samples were washed with PBS 1x, 0.3% Triton x-100 for 1h and mounted in Vectashield containing DAPI or TRITC-Phalloidin (Vector Laboratories). A 120-mm deep spacer (Secure-Seal^TM^ from Sigma-Aldrich) was placed in between the glass slide and the coverslip to preserve morphology of the tissues.

The actin cytoskeleton was labelled using the Rhodamine-Phalloidin probe from Invitrogen (1/500, 2h incubation together with secondary antibodies). Alternatively, samples were mounted in Vectashield hardset medium complemented with TRITC-Phalloidin (Vector Laboratories). Nuclei were labelled with DAPI included in Vectashield mounting medium, when necessary.

Primary antibodies used were: chicken anti β-Gal (GTX77365, 1/500-1/1000, Genetex Inc), mouse anti β-Gal (40-1a, 1/100, DSHB). Chicken anti β-Gal revealed a more intense signal than the mouse one and proved more efficient to reveal basal filopodia-like protrusions. We further used rat anti-ECad (DCAD2, 1/100, DSHB); mouse anti-Dlg1 (4F3, 1/100, DSHB); mouse anti-a-Spectrin (3A9, 1/100, DSHB).

Secondary antibodies used were: anti-mouse Alexa555 (1/200), anti-rabbit Alexa555 and Alexa647 (1/200), anti-rat Alexa555 (1/200) and anti-chicken 649 (1/200). Secondary antibodies were purchased from Invitrogen.

Images were acquired using inverted LSM880 or LSM900 Zeiss laser scanning confocal microscopes fitted with Airyscan modules.

### Live imaging

Dissection tools were cleaned with ethanol before dissection. Leg or wing discs were dissected at white pupal stage + 2/3h in Schneider’s insect medium (Sigma-Aldrich) supplemented with 15% foetal calf serum and 0.5% penicillin−streptomycin as well as 20-hydroxyecdysone (Sigma-Aldrich, H5142,1/100). Leg discs were transferred on a glass slide in 13.5 μL of this medium confined in a 120-μm-deep double-sided adhesive spacer (Secure-Seal^TM^ from Sigma-Aldrich). A glass coverslip was then placed on top of the spacer. Halocarbon oil was added on the sides of the spacer to prevent dehydration.

Imaging was performed using inverted laser scanning confocal microscopes (LSM880 or LSM900 from Zeiss) with C-Apo ×40/1.2 Water Autocorr or Plan-Apochromat x63/1.2 Water objectives from Zeiss.

The Airyscan system was used in most of the recorded movies.

### Laser ablations

For laser ablations, tissues were prepared as described for live imaging using laser scanning microscopy. High quality coverslips (n_1.5H, Marienfeld) were used. Experiments were carried out on both leg and wing prepupal discs. Laser dissection experiments were performed on a Zeiss LSM880 laser scanning microscope fitted with a pulsed DPSS laser (532 nm, pulse length 1.5 ns, repetition rate up to 1 kHz, 3.5 mJ/pulse) steered by a galvanometer-based laser scanning device (DPSS-532 and UGA-42, from Rapp OptoElectronic, Hamburg, Germany). The laser beam was focused through a water-immersion lens of high numerical aperture (Plan-Apochromat 63x from Zeiss). Experiments were performed using a numerical 2x zoom.

Laser intensity for ablations was set at 100% and one or two ablation lines were drawn according to the area to ablate. Also, Z-stacks of the clones were configured to cut through clones’ depth. For laser ablation below the apical surface, ablation lines were 100 pixels long and configured with 100 runs. For this experiment Snail clones expressed a uas::tdTomato in a sqh-eGFP^[KI]^ background. Of note, 532nm laser induces RFP bleaching, therefore a two-colours stack was taken before the experiment to stage the clone and identify the region of interest. Ablation was then run on the green channel only.

For ablations in the filopodia area, ablation lines were 10 to 30 pixels long and configured with 20 to 50 runs. Ablation lines were drawn just below the basal membrane, but not touching it. In these experiments Snail clones expressed either uas::Src1-89-GFP or uas::2xeGFP in a sqh-eGFP^[KI]^ background. Kymographs were obtained by drawing a line on the interest area and using the “reslice” tool in ImageJ. Analyses were based on the quantification of the maximal recoil after the cut.

Clones in which Snail was replaced by a lacZ transgene were taken as control clones for both apical and basal photoablations. Photoablations were performed on clones obtained in both the leg and wing prepupal (0-3h after puparium formation), as clones behave similarly in both tissues.

### Optogenetics

For optogenetics experiments, the uas::PA-Cdc42^[CA]^ construct was crossed to a recombinant line carrying the ap::Gal4^[md544]^ driver and the E-Cad-3mKate2^[KI]^. Crosses were maintained in the dark, and flies were raised at 25°C. Tissues were dissected as indicated above, but with minimal light exposure.

Movies of the distal leg were acquired based on the red channel only (using 561nm excitation, a waveline that does not activate PA-Cdc42^[CA]^). Images were taken in a single z plane every 0.5sec.

After 20sec of recording a 2.2*2.2µm ROI (region of interest) placed on the most basal part of the cells was used to activate PA-Cdc42^[CA]^ using a single pulse of the 488nm laserline set at 100%. Maximal apical-to-basal displacement of E-Cadherin was analysed on kymographs.

### Quantifications

#### Intensity of fluorescent molecules

Intensity of fluorescent molecules in EMT clones was quantified in EMT clone’s sagittal views with Image J, either by drawing a line of dots. For basal myosin-GFP or Dg-GFP intensity, a line with a width of 10px was drawn by following the basal surface of the clone in one z-plane. This measurement was performed in three different z-planes for each EMT clone and then averaged to obtain a homogenised intensity value for each clone. The same quantification method was performed in neighbouring wild-type cells, the internal control. Basal myosin or Dg intensity in EMT clones was then normalised by their associated internal control. A similar strategy was used for apical quantification of Dlg.

For apical E-Cadherin-GFP or Crumbs signal, a circle corresponding to the ECadherin or Crumbs signal was drawn with Image J. For each clone, the signal was measured in three different z-planes in which the dots were evident to contour. The three measures per clone were then averaged. The same quantification method was performed in neighbouring wild-type cells, the internal control. Then, for each clone the E-Cadherin or Crumbs intensity was normalized by their associated internal control. A similar strategy was used for basal Talin quantification.

#### Number & length of filopodia-like protrusions

Number & length of filopodia-like protrusions were quantified with Image J. Filopodia has been measured in maximum projection for each EMT clone in a sagittal view. A segmented line was drawn from the filopodia base to the filopodia tip.

#### Analysis of apical surface deformation

Apical surface deformations induced by EMT clones were quantified with the line tool of ImageJ. Z-stacks of EMT clones were imaged and the deformation depth measurement was done in the z-plane in which the deformation was the higher. As the epithelium’s height can vary according to the leg’s area, the % of epithelium height pulled was instead quantified by dividing the deformation depth in microns with the wildtype epithelium’s height (also in microns) next to the clones.

#### Stage repartition following alteration of filopodia-like protrusions

Stage repartition following impairment of filopodia-like protrusions was based on the clone’s shape and subsequent integration or not within the epithelium. All the leg’s EMT clones were analysed and distributed in two main categories: fully integrated clones (early EMT stages) or partially out clones (*i.e.* undergoing cell exit). To do so, each z-plane of the stack was analysed in Image J and the orthogonal clones were visualized with Zen OrthoSlicer tool.

### Statistical analysis

Statistics were performed in Prism. n and p values are indicated in figure legends. Box plot were generated in Prism and represent the median, with minimal and maximal range. Violin plots and dots repartition graphs (in which red bars indicate the mean +/- Standard error of the mean, SEM) were also generated in Prism.

The normality of data sets was determined using Prism 8 (GraphPad). For data sets that follow a normal law, a two-tailed unpaired Student’s t-test was used to assess the significance. Otherwise, a Mann and Whitney test was performed. We used T-test for the analysis of candidates from the genetic screen.

Significance is denoted as follows according to the p value: ****p<0.0001; ***p<0.0005; **p<0.001; *p<0.05; NS p>=0.05 (not significant).

Box plot were generated with Prism.

---

**Figure S1.**
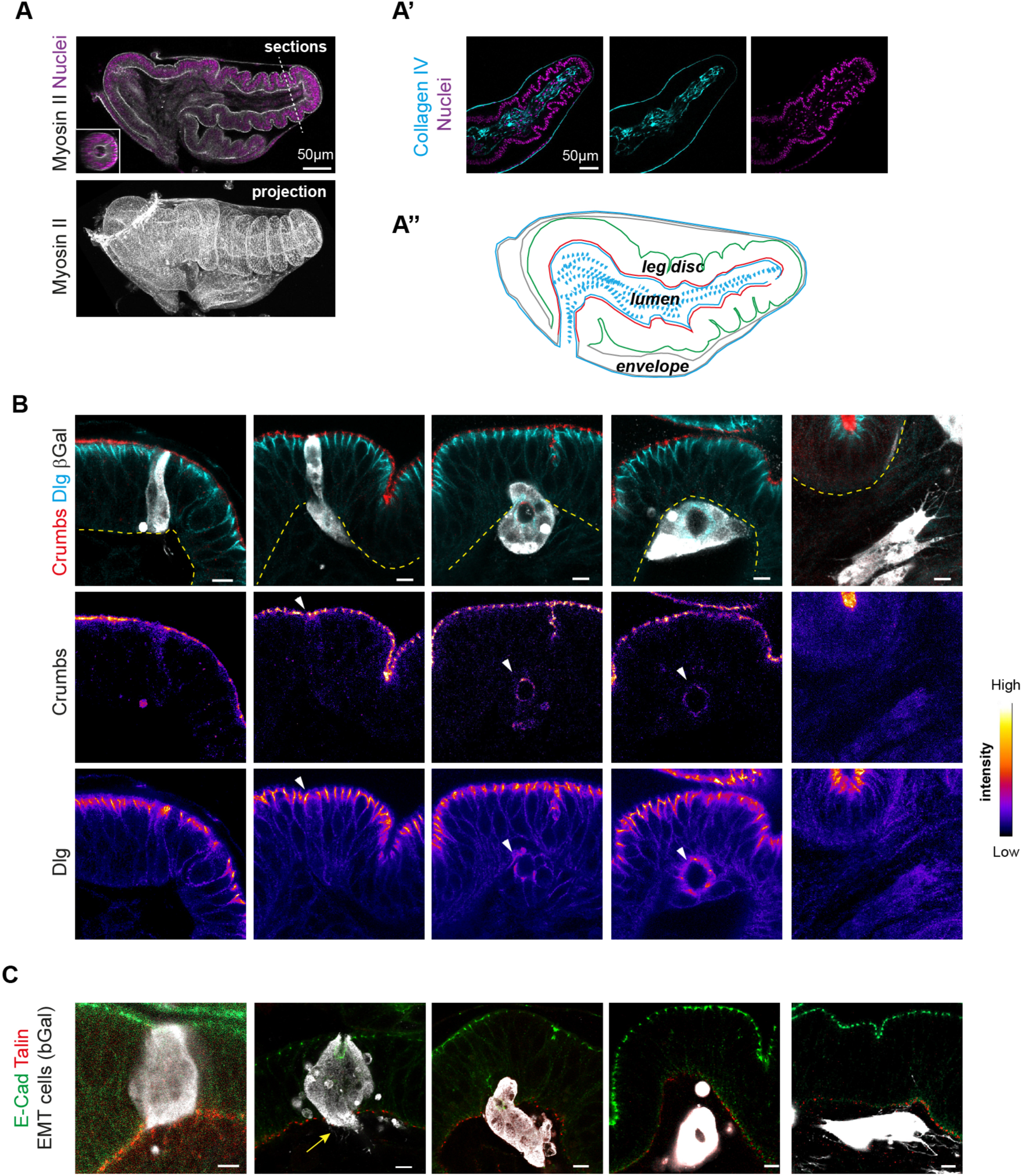
Epithelial-to-Mesenchymal transition dynamics in the *Drosophila* leg. Related to Figure 1. **(A-A’’)** Illustration of the *Drosophila* prepupal leg (white pupa +2 hours after puparium formation). (A) Transversal confocal section along the proximodistal axis (top) and 3D reconstruction (bottom) showing myosin II-GFP distribution and the overall tissue shape. Nuclei are shown in magenta in the transverse section. Inset corresponds to a section at the level of the dash line. The *Drosophila* prepupal leg is a monolayer epithelium forming a cylindrical tissue around a central lumen. The epithelium is closed at the distal extremity (on the right), but open on the proximo-ventral part. The leg epithelium is surrounded by an envelope (known as the peripodial membrane). Dorsal is up. (A’) ECM is revealed by Collagen IV-GFP staining. ECM accumulates within the lumen, where muscle precursors and cells of the nervous systems are found. It also accumulates at the base of the envelope. (A’’) Schematization (transversal section representation) of the prepupal leg. **(B, C)** Timeline of Snail-induced EMT in the fly leg. Images correspond to images shown in Figure 1. The white channel (β-Gal staining) corresponds to the EMT cells, and is schematized by the dashed white lines in Figure 1. In (B), single channels for Crumbs and Dlg are shown using an intensity color-code. In (C), the yellow arrow points at the basal of the Snail clone invading the underlying luminal compartment. Scale bars: 5µm unless indicated.

**Figure S2.**
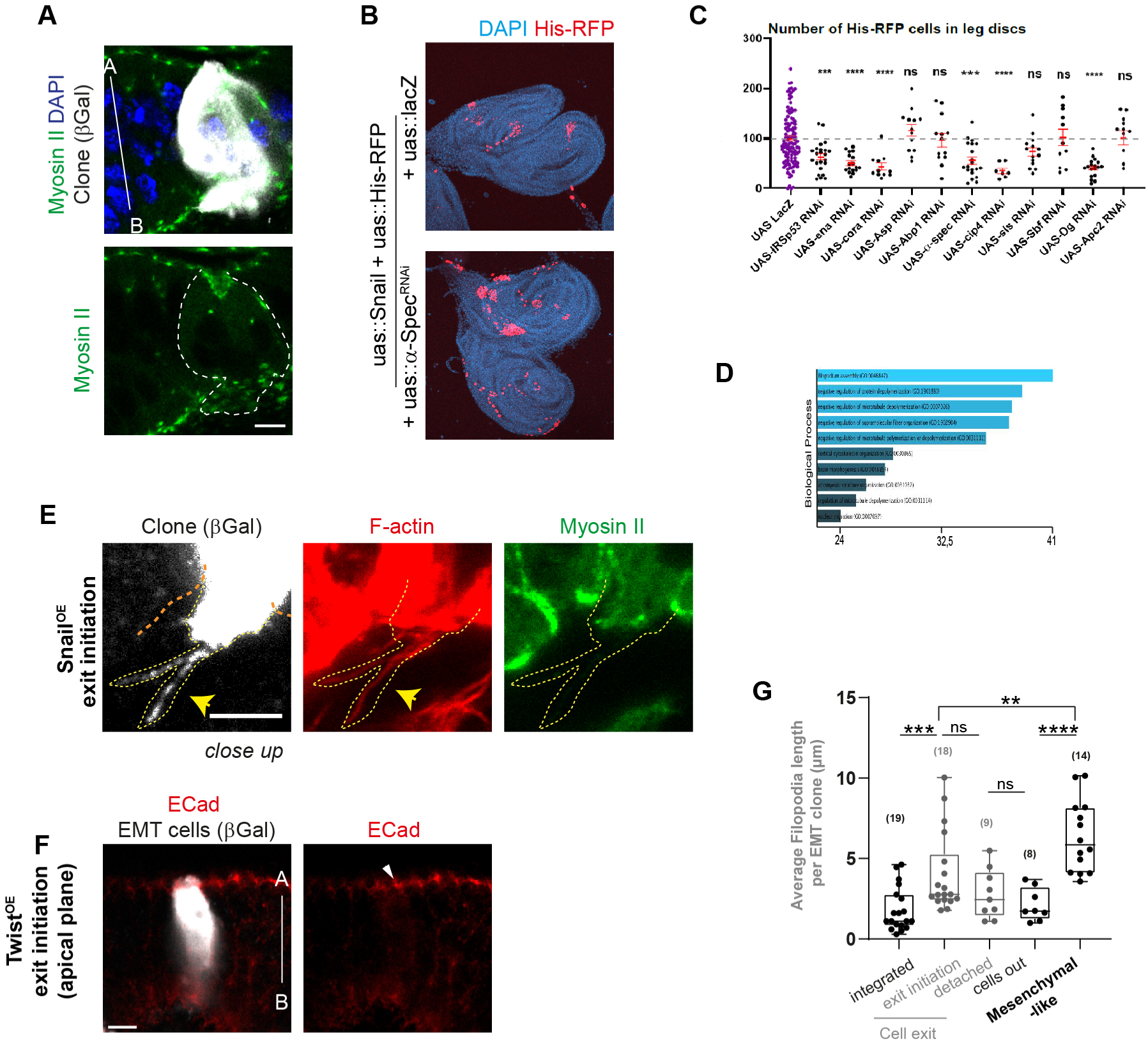
EMT cells generate basal filopodia-like protrusion during cell exit. Related to Figure 2. **(A)** A Snail-expressing clone at the exit initiation stage (still attached apically and invading basally the underlying luminal compartment) stained for myosin II-GFP (green). Note that the apical surface deformation is not associated with strong apico-basal myosin II structures within the clone (n=17/22). **(B)** An example from the genetic screen, where the overall disc morphology is shown by DAPI staining. His-RFP labels EMT cells. **(C)** Quantification of cell numbers (normalized by the mean of the lacZ control) following RNAi expression in clones without Snail expression (secondary screen). Each dot corresponds to the average number of cells in a given leg. The red bars indicate the mean and standard error of the mean (SEM). **(D)** Graph highlighting Gene Ontology associated biological functions of the candidates from the genetic screen. **(E)** Close up view showing F-actin staining and Myosin II within Snail-induced EMT cells. Arrows indicate filopodia-like structures enriched in F-actin. **(F)** Cell exit induced by a distinct EMT regulator, Twist. This image focuses on the apical plane of the clone shown in Fig.2C (which was focusing on the basal side). Note that Twist-expressing cells at the exit initiation stage maintain apical E-Cadherin staining (arrowhead), similar to Snail-expressing cells (Fig.1C). **(G)** Quantification of filopodia length in Snail-expressing cells. n are indicated in brackets. Scale bars: 5µm. In box plots, the black bar and whiskers indicate the median and the maximal range. Statistical tests: Mann and Whitney in C, G. p-values: *, <0.01; **, <0.001; ***, <0.0005; ****, <0.0001; ns, non significant.

**Figure S3.**
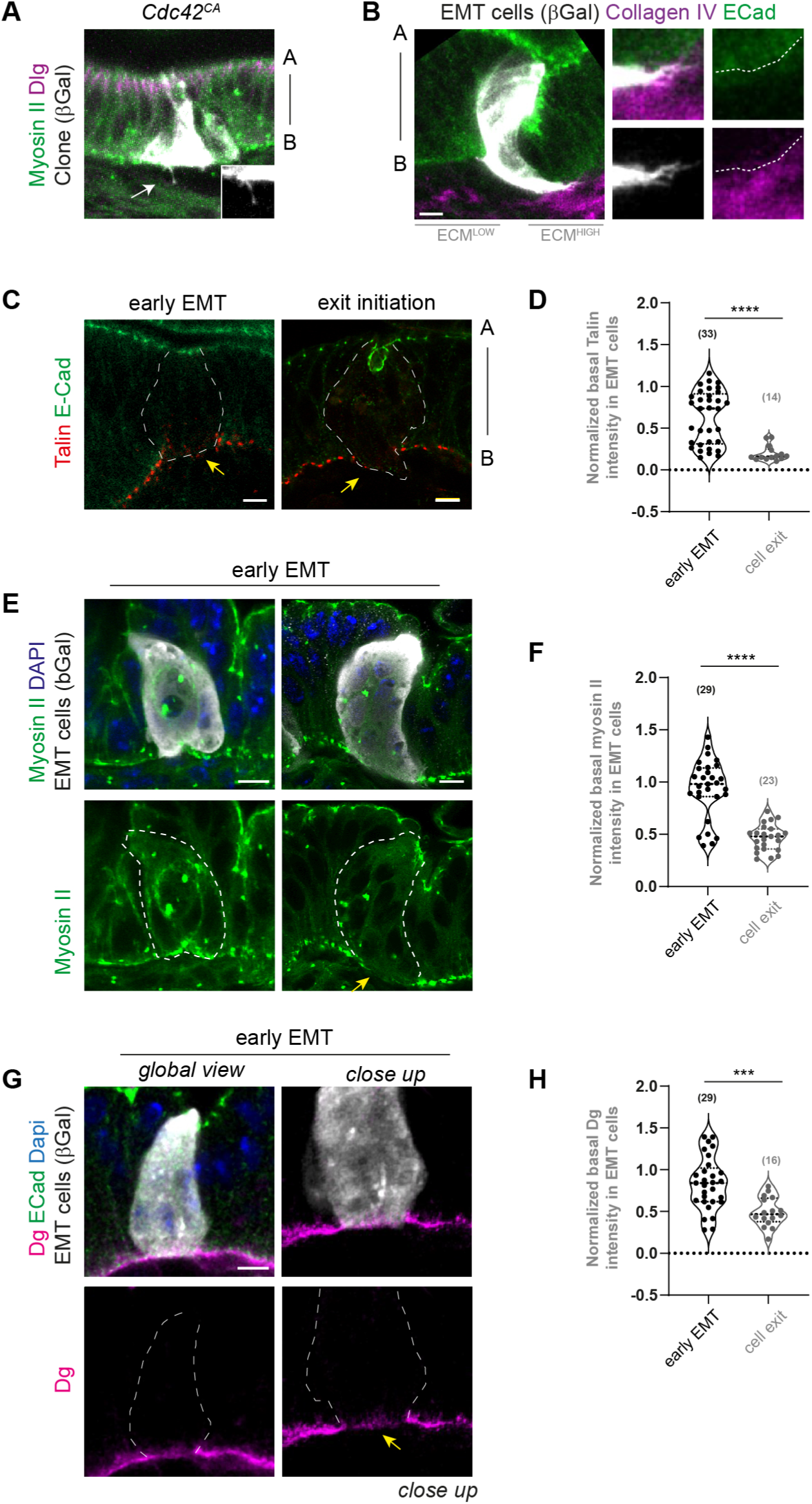
Basal dynamics in Snail-induced EMT cells. Related to Figure 3. (A) A clone of cells expressing the constitutive active form of the Cdc42 filopodia regulator in the fly leg. Note that Cdc42^CA^ is sufficient to induce filopodia formation (arrow, see inset). (B) A Snail-expressing clone at the exit initiation stage which has started to invade the underlying luminal compartment. Inset shows filopodia-like structures at the extremity of the clone. The ECM is shown in magenta. Regions with low and high levels of ECM are indicated. Note that the clone tip is in close vicinity with the region of high ECM intensity. The dash lines indicate the basal of the epithelium. **(C-H)** Images of Snail-induced EMT clones (C, E, G) and corresponding quantifications represented as violin plots (D, F, H) of basal markers (Talin, basal myosin II, Dg) at the indicated stages. n analyzed are indicated in brackets. Clones outlines are indicated by white dash lines. Note that clones shown in C correspond to images shown in Fig.S1. Yellow arrows point at basal depletion of the indicated markers. Scale bars: 5µm. Statistical tests: Mann and Whitney in D, F, H. p-values: ***, <0.0005; ****, <0.0001.

**Figure S4.**
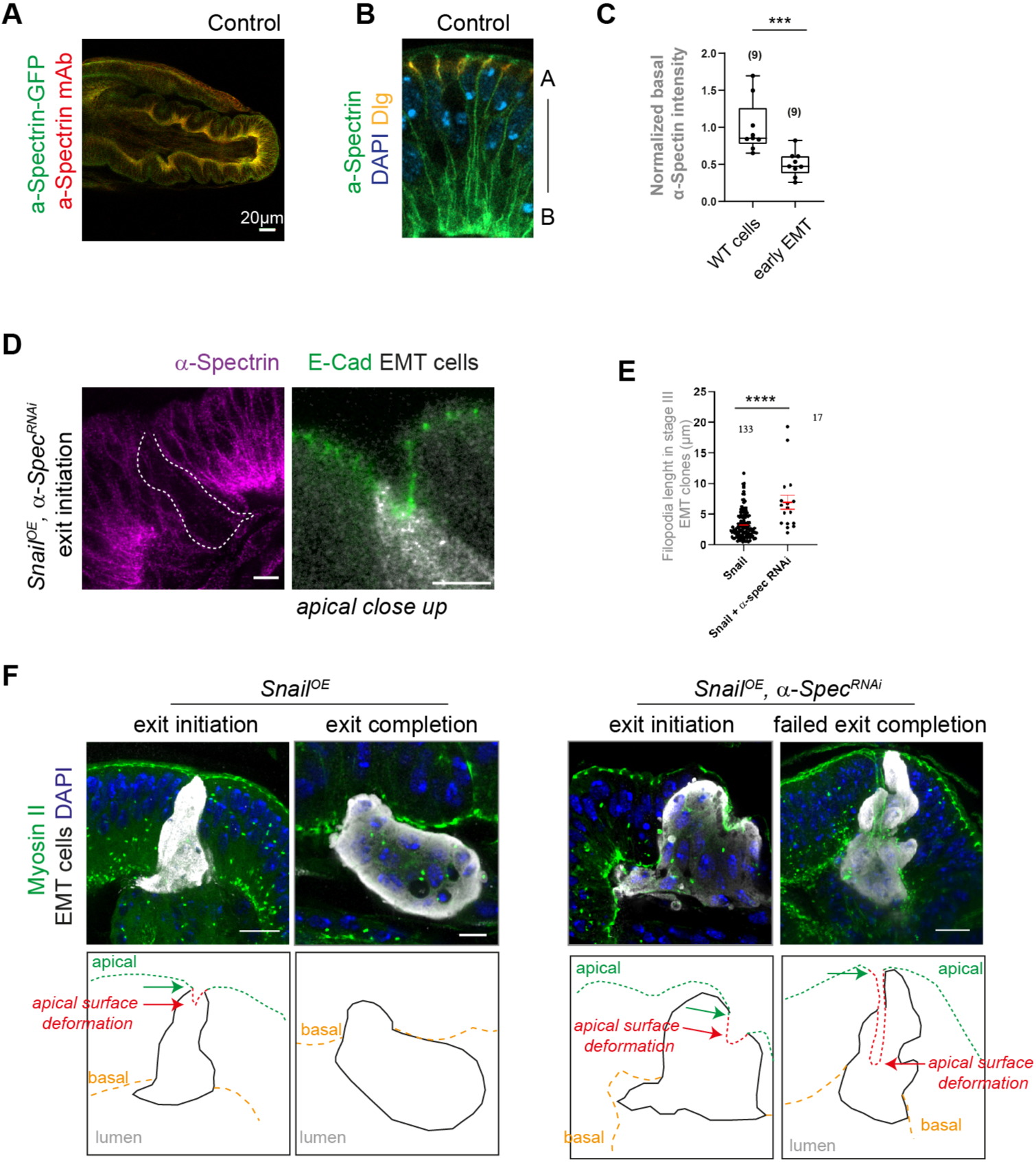
Analysis of the lateral α-Spectrin cytoskeleton. Related to Figure 4 (A) Global view showing α-Spectrin distribution in the *Drosophila* pupal leg. α-Spectrin is detected with an antibody (red) and a GFP insertion at the endogenous locus (green). (B) Close up view showing α-Spectrin localization up to the apical domain which is revealed by Dlg staining (orange). (C) Quantification of normalized basal α-Spectrin-GFP intensity in wild-type (WT) and Snail cells at the early EMT stage. (D) Immunostaining revealing nearly complete α-Spectrin depletion in the Snail clone (outlined in white) following co-expression of the α-Spectrin^RNAi^ (left). Right: close up view of the apical domain of the clone, showing the normal distribution of E-Cadherin despite α-Spectrin depletion at the exit initiation stage. These images correspond to Fig.4B. (E) Quantification of filopodia length in Snail^OE^, α-Spectrin^RNAi^ cells. (F) Transverse views of EMT cells either control or co-expressing α-*Spectrin^RNAi^*shown in Fig.4D. The clones are detected thanks to the β-Gal staining (white). Schematization of the clones is shown on the bottom line. Red arrows point at maximal apical surface deformation, while green arrows indicate the base of the apical surface. Scale bars: 5µm unless indicated. Statistical tests: Mann and Whitney in C, E. p-values: ***, <0.0005; ****, <0.0001;.

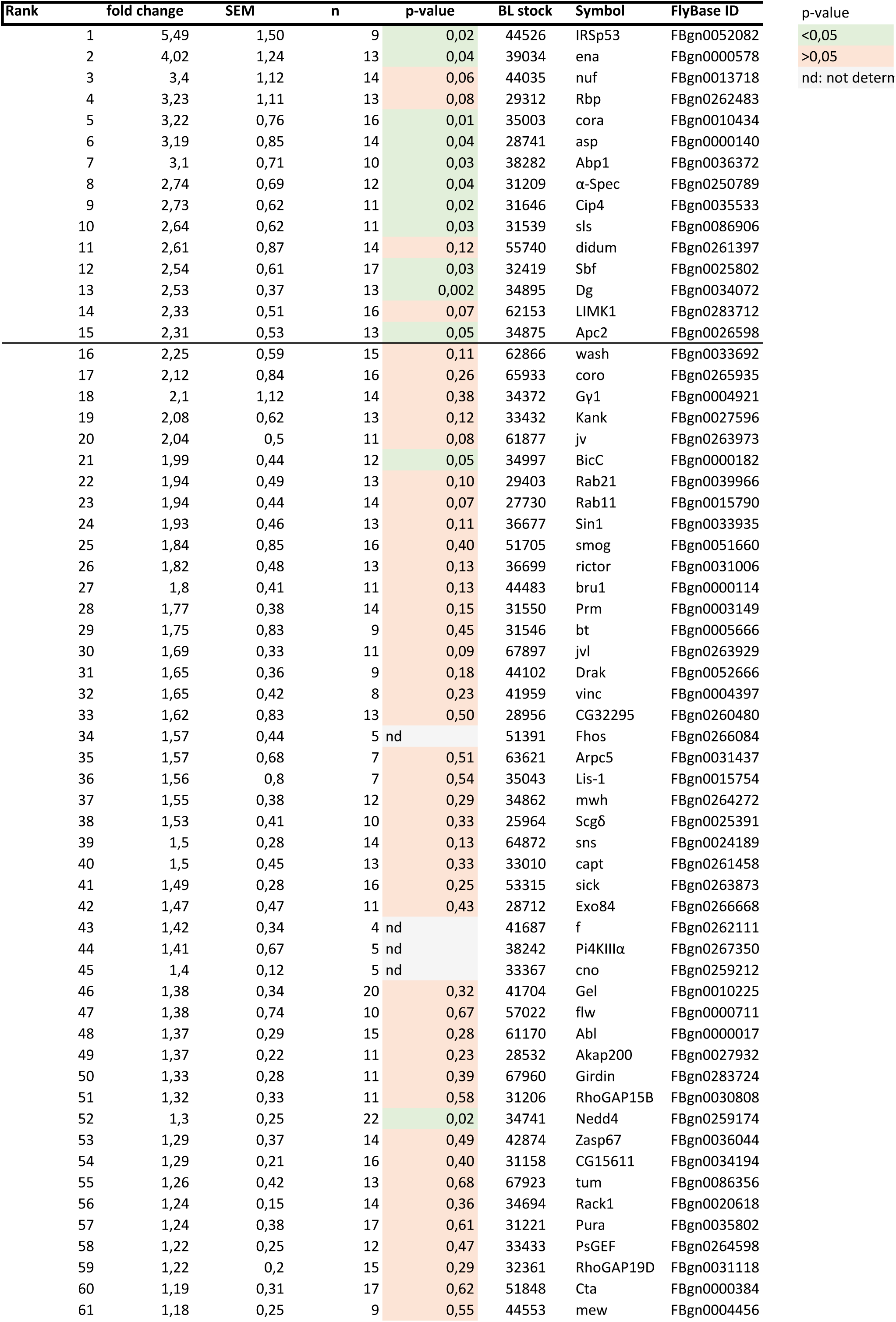
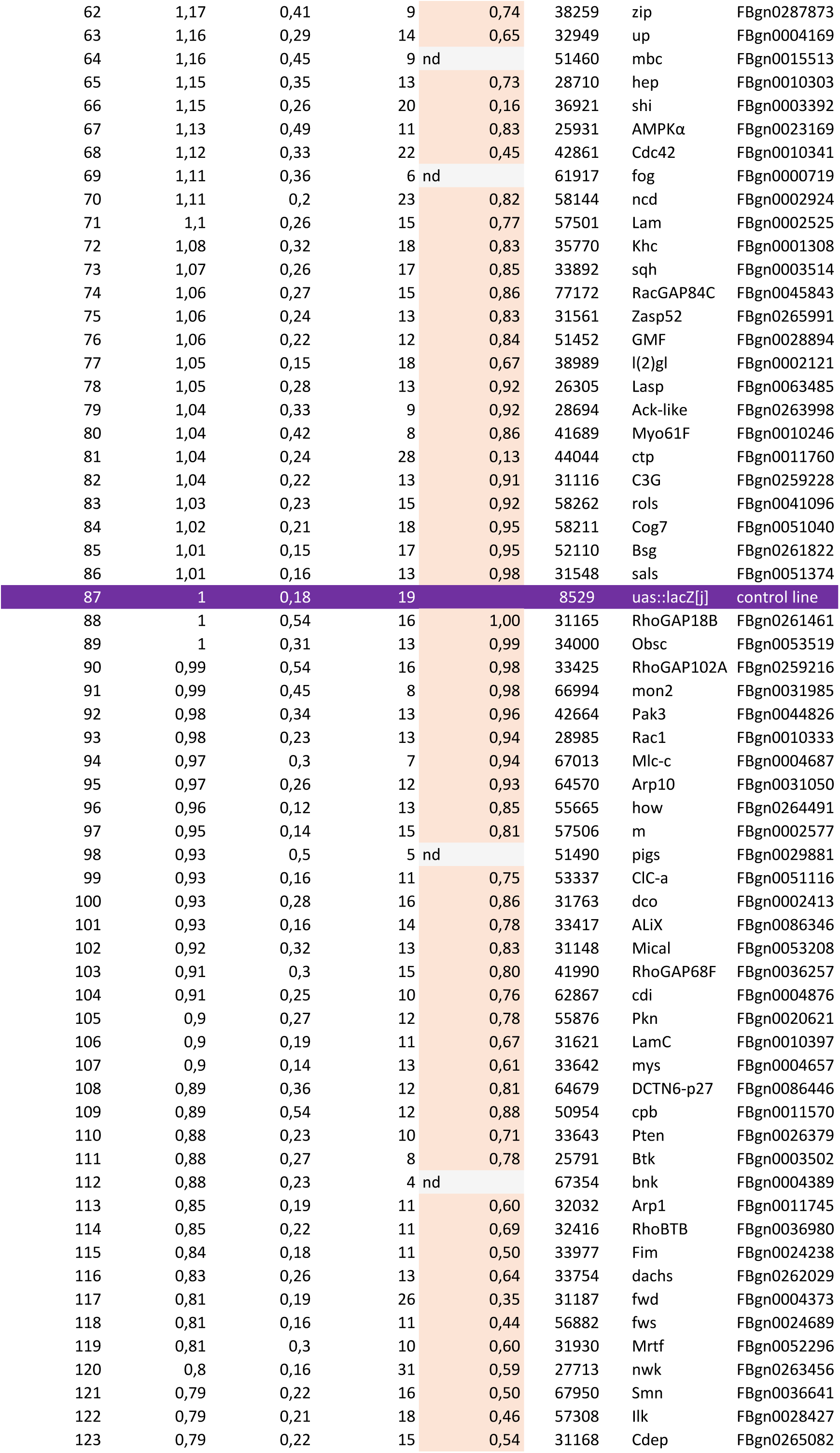
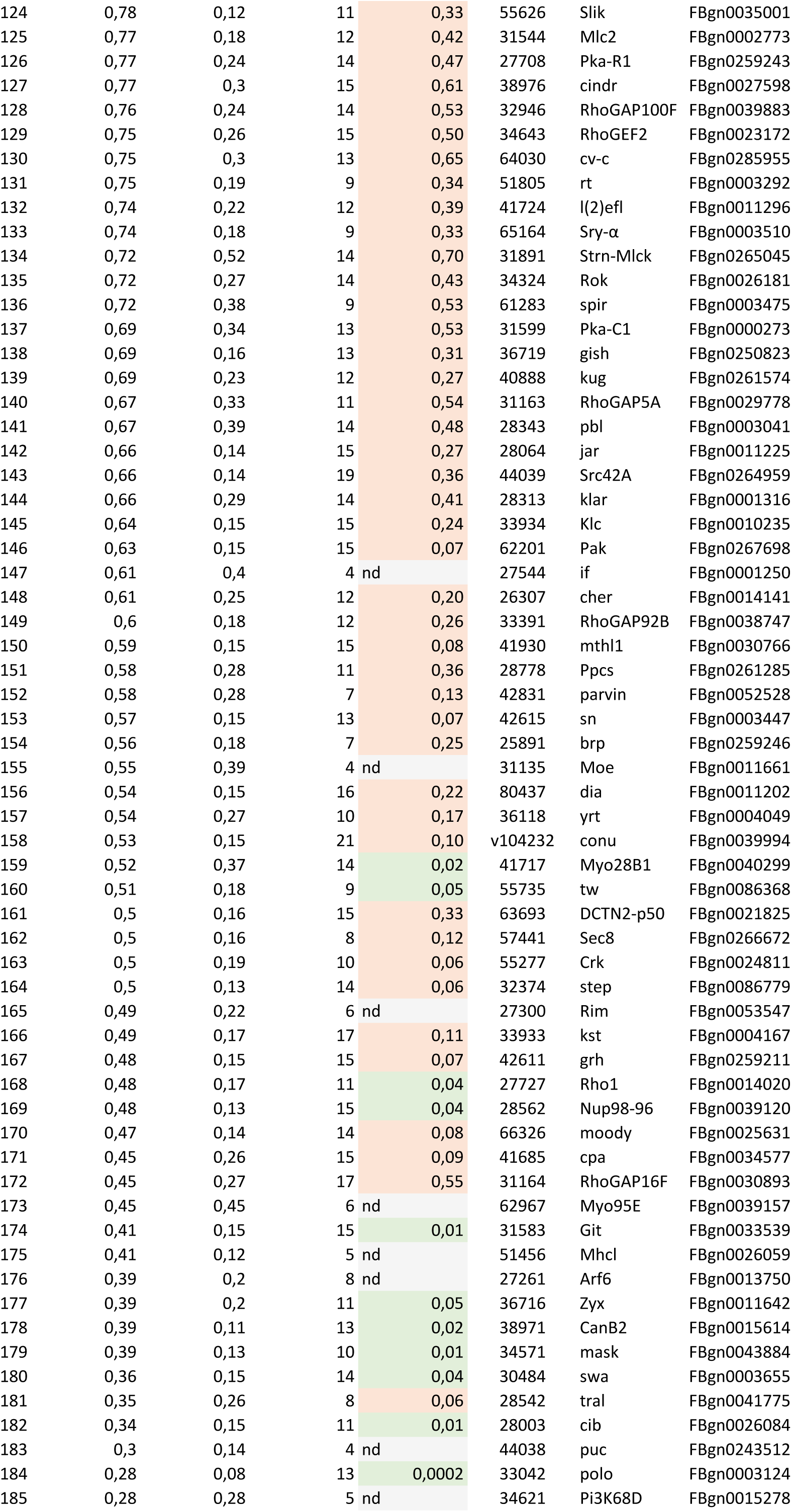
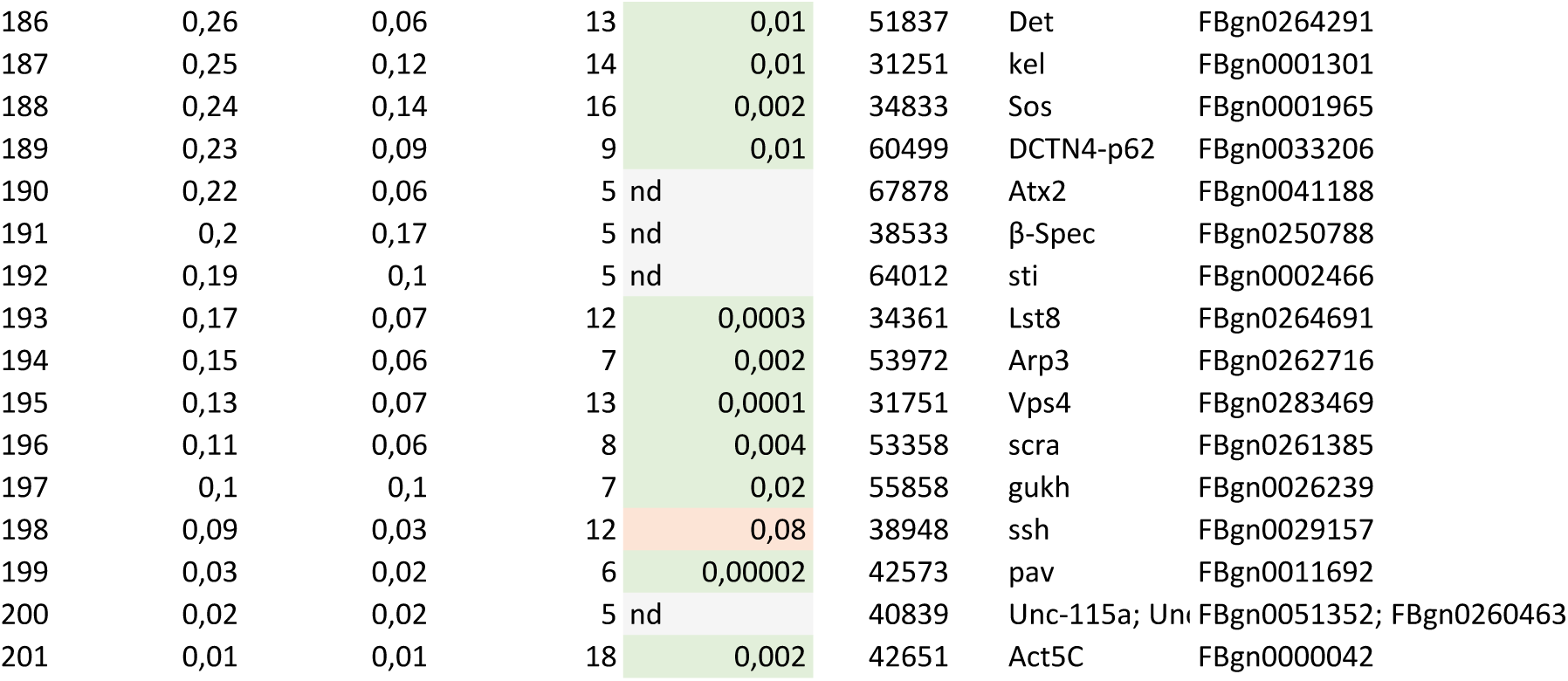

